# The effects of *E. granulosus* PSCs proteins on host immune cells were elucidated through Transcriptomic and Metabolomic analyses

**DOI:** 10.1101/2025.05.06.652359

**Authors:** Kalibixiati Aimulajiang, Xia Chen, Mayire Aizezi, Tanfang Zhou, Renyong Lin, Hao Wen

## Abstract

**Background:** *Echinococcus granulosus* (*E. granulosus*), a causative agent of zoonotic diseases, is widely prevalent across most regions and primarily affects vital organs such as the liver and lungs. The parasite secretes immunomodulatory substances, and its protoscolex proteins (PSCs) have been identified as potential diagnostic antigens and vaccine candidates. Although transcriptomics and metabolomics approaches have been employed to investigate echinococcosis, the impact of PSCs on the functionality of host immune cells and the expression of immunomodulatory-related genes remains largely unexplored.

**Methods:** In this study, immunofluorescence analysis was employed to determine the binding of PSCs to lymphocytes. The CCK-8 assay was utilized to evaluate the impact of varying concentrations of PSCs on cell viability, while ELISA was performed to quantify the levels of cytokines such as IL-9 and IL-4. Furthermore, by integrating LC-MS/MS-based metabolomics and RNA-Seq-based transcriptomics technologies, a systematic analysis was conducted to investigate the metabolic reprogramming and differential gene expression in lymphocytes following PSCs treatment, thereby elucidating the molecular mechanisms underlying its immune regulatory effects.

**Results:** PSCs proteins inhibited lymphocyte proliferation in a concentration-dependent manner, promoted IL-9 secretion at low concentrations, enhanced IL-4 expression at medium concentrations, and continuously up-regulated TGF-β with increasing concentrations. Transcriptomic analysis identified 1,840 differentially expressed genes (991 up-regulated and 849 down-regulated), and KEGG enrichment analysis highlighted metabolic pathways, including phenylalanine metabolism. Gene Ontology (GO) analysis revealed that the functions of these genes were primarily associated with immunometabolic reprogramming and regulation of the intestinal barrier. Metabonomic profiling detected 12 significantly altered metabolites (8 up-regulated and 4 down-regulated). These findings suggest that PSCs reshape the host immune microenvironment via a metabolic-gene network. This study systematically elucidates how PSCs regulate immune cell function and metabolic homeostasis through multiple targets, providing novel insights into the immune escape mechanisms of parasites.

**Conclusion:** PSCs can induce host immune metabolic reprogramming, characterized by the activation of pyrimidine metabolism and the inhibition of bilirubin-mediated oxidative stress, through concentration-dependent regulation of Th9 cytokines (IL-9, IL-4/TGF-β) and suppression of the PI3K-Akt signaling pathway. This process remodels the immune tolerance microenvironment. The aim is to elucidate the molecular mechanisms underlying parasite escape via multi-dimensional immunometabolic regulation, thereby providing a novel direction for the targeted treatment of echinococcosis.

**Author summary:** *E. granulosus* is a zoonotic disease that can cause space-occupying lesions in the liver, lung and other organs, seriously threaten human health and cause economic losses in animal husbandry. It evades immune attack through a variety of mechanisms.This study investigates the regulatory effects of *E. granulosus* PSCs on host immune cell function, with a focus on their concentration-dependent influence on Th9 cell differentiation and cytokine expression, including IL-9. Through integrative analyses of transcriptomics and metabolomics, we elucidate the molecular mechanisms underlying immunometabolic reprogramming induced by PSCs. Our findings demonstrate that at low concentrations, PSCs promote the secretion of IL-9 associated with Th9 cells, upregulate IL-4/TGF-β signaling, activate pyrimidine metabolism (leading to thymidine accumulation), enhance bile secretion pathways, and increase bilirubin levels to suppress oxidative stress and T-cell function, thereby establishing an immune-tolerant microenvironment. This research provides the first comprehensive illustration of immune microenvironment remodeling mediated by PSCs via multi-omics network regulation, offering novel insights into the treatment of cystic echinococcosis and host-parasite interactions.

## Introduction

*E. granulosus*, a zoonotic disease, is caused by infection with the parasite *E. granulosus* is caused by infection with the parasite *E. granulosus*, and it remains endemic in most countries and regions, although it has been eliminated in certain countries[1]. In endemic areas, the annual incidence of CE is less than 1 to 200 per 100,000 individuals. CE infection primarily affects and causes damage or dysfunction in target organs, particularly the liver (70%) and lungs (20%). Other affected organs include the brain, spleen, kidneys, and heart[1–3].Worms are known to release ESPs to the host, which possess potent immunomodulatory properties and exert a wide range of effects on host immunity[4]. In addition, the protoscolex protein of *E. granulosus* can serve as a diagnostic antigen for detecting serum antibodies in infected patients, exhibiting remarkable immunogenicity [5]. Another study found that, The combination of Psc-derived recombinant proteins rEgTIM with rEgANXB3, rEgADK1with rEgEPC1, and rEgFABP with rEgA31 were resistant to *E. granulosus* infection in dogs[6].Crude protein from PSCs has also been used as a vaccine for immunization of the definitive host[6].The viability of PSCs is a crucial determinant for the developmental competence of CE cysts[7] and is influenced by the host species and immune responses[8]. However, the exact biological function and immunological characters of PSCs protein are not clear.

The application of transcriptomics and metabolomics has been utilized in the investigation of immunopathological and cellular functional aspects related to echinococcosis[9–11]. Numerous studies in the field of transcriptomic analysis have been dedicated to investigating *E. granulosus* protoscolex, with a particular focus on elucidating the differential expression patterns of ESPs[12], alternative splicing events[13], and genes involved in the bi-directional transformation towards either adult or cystic forms[14, 15].The application of metabolomics has revealed a significant association between phenylalanine metabolism and the pathogenesis of *cystic echinococcosis*[16], while also confirming that infection with *E. granulosus* protoscoleces induces metabolic reprogramming in B cells of mice[9]. However, these analyses did not investigate the impact of *E. granulosus* protoscoleces proteins on host immune cell functionality or the differential gene expression associated with immunomodulation.

T cells have been implicated in modulating the immune milieu within the hepatic microenvironment of patients afflicted with CE and mice infected with *E. granulosus* sensu stricto; Among these immunological cells, CD4 T cells were identified as the predominant subset, and their participation was intricately associated with both cyst viability and establishment[17]. Recent studies have shown that IL-9 can be secreted by a specialized T cell population called Th9 cells, which mediate anti-parasite immunity by producing IL-9[18]. Notably, both systemic TGF-β/IL-9 levels and hepatic IL-9/IL-9 receptor expression are significantly heightened in individuals affected by cystic echinococcosis of the liver.[19]. IL-9 concentrations demonstrate a direct correlation with hepatic inflammatory severity[19]. Our prior investigations further revealed that *E. granulosus* cyst fluid modulates Th9 cell differentiation and its cytokine secretion in vitro, elucidating mechanisms underlying host-parasite interactions[20] .

This study sought to characterize the immunoregulatory effects of *E. granulosus* PSCs on murine splenic lymphocytes, with a focus on their capacity to drive Th9 cell polarization and IL-9 secretion. Integrated transcriptomic and metabolomic profiling validated the systemic impact of PSCs proteins on lymphocyte activity. These findings elucidate novel mechanisms of host immune modulation by parasitic agents, providing a foundation for targeted therapeutic interventions against cystic echinococcosis.

## Materials and methods

### Animals

Female Balb/c mice aged 6-8 weeks were procured from Xinjiang Medical University and housed in a meticulously maintained facility within the same institution, strictly adhering to standardized protocols. The animal study was conducted in accordance with the guidelines set forth by the Animal Ethics Committee (approval number: IACUC-20210301-01) at Xinjiang Medical University.

### Acquisition of PSCs and preparation of their proteins

Sheep livers infected with *E. granulosus* were obtained from local abattoirs and sterilized by spraying alcohol. First, the cyst fluid of *E. granulosus* was aseptically extracted, and protoscolex was naturally precipitated. Next, the supernatant was discarded and the natural sediment was washed with sterile saline, and collected after five repetitions. Digestion was performed with 1% pepsin (Solarbio, Beijing, China) in a bath solution at 37℃ for 30 min and was removed and mixed every 5 min. Finally, the above protoscolex solution was centrifuged and the supernatant was discarded, stained with 1% trypan blue (Solarbio, Beijing, China) and counted using a microscope.

The fresh protoscolex was put into 2ml tube. Firstly, 500ul calcium-free magnesium PBS solution and two 4mm steel balls were added. They were then put into a precooled tissue grinder for 6 min. Following homogenization, the sample was subjected to overnight agitation at 4°C. Subsequent centrifugation (13,000 × g, 10 min, 4°C) yielded a clarified supernatant, which was aseptically filtered (0.22 μm membrane) and archived at -20°C. Protein quantification of protoscoleces lysates was performed using a BCA assay kit (Solarbio, Beijing, China).

### Isolation of mouse spleen lymphocytes

Following aseptic extraction, the spleen was transferred to a sterile culture dish immersed in ice-cold PBS. The tissue underwent mechanical disruption via frosted slide grinding, followed by nylon mesh filtration (70 μm pore size) to obtain a single-cell suspension. Lymphocyte isolation was performed according to the manufacturer’s protocol (Mouse Spleen Lymphocyte Separation Kit, Solarbio, China). Cell viability and density were quantified using trypan blue exclusion prior to downstream applications.

### Immunofluorescence

Murine splenic lymphocytes (1×10⁶ cells/mL) were seeded in 24-well plates and treated with 100 μg/mL PSCs, with untreated wells serving as controls. Following a 2-hour incubation at 37°C with 5% CO_2_, cells were harvested, washed twice in PBS, and cytospun onto poly-L-lysine-coated slides for 15-min adhesion. After discarding supernatants, samples were fixed with 4% paraformaldehyde (RT, 15 min) and blocked with 5% BSA. Primary immunostaining employed rat polyantiserum (1:500 dilution), paralleled by negative controls using non-immune rat serum (1:500). Secondary detection utilized DyLight594-conjugated goat anti-rat IgG (1:300, BA1141, BOSTER) under light-protected 37°C incubation for 1 h. Nuclear counterstaining was performed with DAPI (Servicebio) in dark conditions (RT, 30 min). Fluorescent signals were visualized using a Nikon A1R confocal system with NIS-Elements imaging software.

### Cell viability assay

Splenic lymphocytes from mice were exposed to escalating PSCs concentrations (25, 50, 100, and 200 µg/mL) for 24 hours at 37°C under 5% CO_2_, with PBS-treated cells as controls. Cellular viability was evaluated using the CCK-8 assay (Beyotime, China). Following treatment, 10 μL of CCK-8 reagent was added to each well, and optical density at 450 nm (OD450) was quantified using a Multiskan™ microplate reader (Thermo Scientific, USA).

### Measurement of cytokines

Murine splenic lymphocytes were treated with PSCs at escalating doses (25-200 µg/mL) versus PBS controls under standard culture conditions (37°C, 24 hr). Cytokine profiles (IL-4, IL-10, TGF-β, IL-9) in supernatants were analyzed using commercial ELISA kits (Lapuda, China) according to the manufacturer’s protocols. Absorbance at 450 nm was quantified with a Multiskan™ microplate reader (Thermo Fisher, USA).

### LC–MS/MS analysis

Liquid chromatography was conducted on a Thermo Scientific™ Ultimate 3000 UHPLC system equipped with an ACQUITY UPLC® HSS T3 column (150 × 2.1 mm, 1.8 μm; Waters) maintained at 40°C. Operational parameters included a flow rate of 0.25 mL/min and an injection volume of 2 μL. Mass spectrometric detection utilized a Q Exactive HF-X Orbitrap platform (Thermo Fisher) controlled by Xcalibur software (v4.3), with full-scan MS data acquired in positive/negative ionization modes. Key ESI conditions were optimized as follows: spray voltage ±3.5 kV (positive/negative); capillary temperature 320°C; sheath gas 50 Arb, auxiliary gas 10 Arb; MS resolution 70,000 (full scan), 17,500 (MS/MS); normalized collision energy 30%.

Raw MS data were converted to mzXML files via ProteoWizard (v3.0.20) and processed through a custom R-based pipeline integrating XCMS algorithms. Metabolite annotation was performed against the BiotreeDB MS² spectral library with a similarity threshold of 0.3. Following log-transformation and unit variance scaling (SIMCA-P+ 16.0.2), orthogonal partial least squares-discriminant analysis (OPLS-DA) was implemented to evaluate metabolic variance, with model robustness validated through seven-fold cross-validation.

### RNA-Seq

Total RNA was extracted using TRIzol® reagent (Invitrogen), and its purity and integrity were assessed via NanoDrop 2000 (Thermo Fisher). Library construction followed the Illumina Stranded mRNA protocols: Poly-A+ RNA selection using magnetic beads, followed by chemical fragmentation (Illumina buffer). First-strand cDNA synthesis employed random hexamers and SuperScript™ II Reverse Transcriptase, with second-strand generation using DNA Pol I and RNase H. Blunt-end repair and adapter ligation (Illumina PE) preceded size selection (400-500 bp) via AMPure XP beads (Beckman Coulter Beverly, CA, USA). Library amplification utilized Illumina primers (15 PCR cycles), followed by quantification on a Bioanalyzer 2100 (Agilent HS DNA Kit). Final sequencing was performed on an Illumina NovaSeq 6000 platform (PANOMIX, Suzhou).

Gene expression quantification was performed using HTSeq (v0.9.1) with raw read counts normalized to FPKM values. Differential expression analysis **was carried out using** DESeq2 (v1.30.0) with stringent thresholds: |log₂FC| > 1 and adjusted P < 0.05. Bidirectional hierarchical clustering of differentially expressed genes (DEGs) was visualized via pheatmap (v1.0.8). Functional annotation of DEGs included GO term enrichment through topGO and KEGG pathway analysis using clusterProfiler (v3.4.4).

### Statistical analysis

Data were derived from three independent experimental replicates and expressed as mean ± SEM. Statistical analyses were performed using one-way ANOVA with post hoc comparisons, considering \**p*\* < 0.05 as statistically significant

## Results

### *E.granulosus* PSCs Inhibit Lymphocyte Proliferation and Modulate IL-9, IL-4, TGF-β Concentration-Dependently

Immunofluorescence analysis validated in vitro binding between PSCs and murine splenic lymphocytes (Fig 1A). PSCs-treated samples exhibited intense red fluorescence at mouse spleen lymphocytes interfaces, with colocalization signals visualized as purple emission. Negative controls displayed no specific fluorescence, confirming PSCs-mediated cellular interactions. Subsequent CCK-8 assays revealed dose-dependent suppression of lymphocyte viability, showing maximal inhibition at intermediate concentrations (100 µg/mL) but attenuated effects at higher doses (200 µg/mL) (Fig 1B).

**Fig 1:**
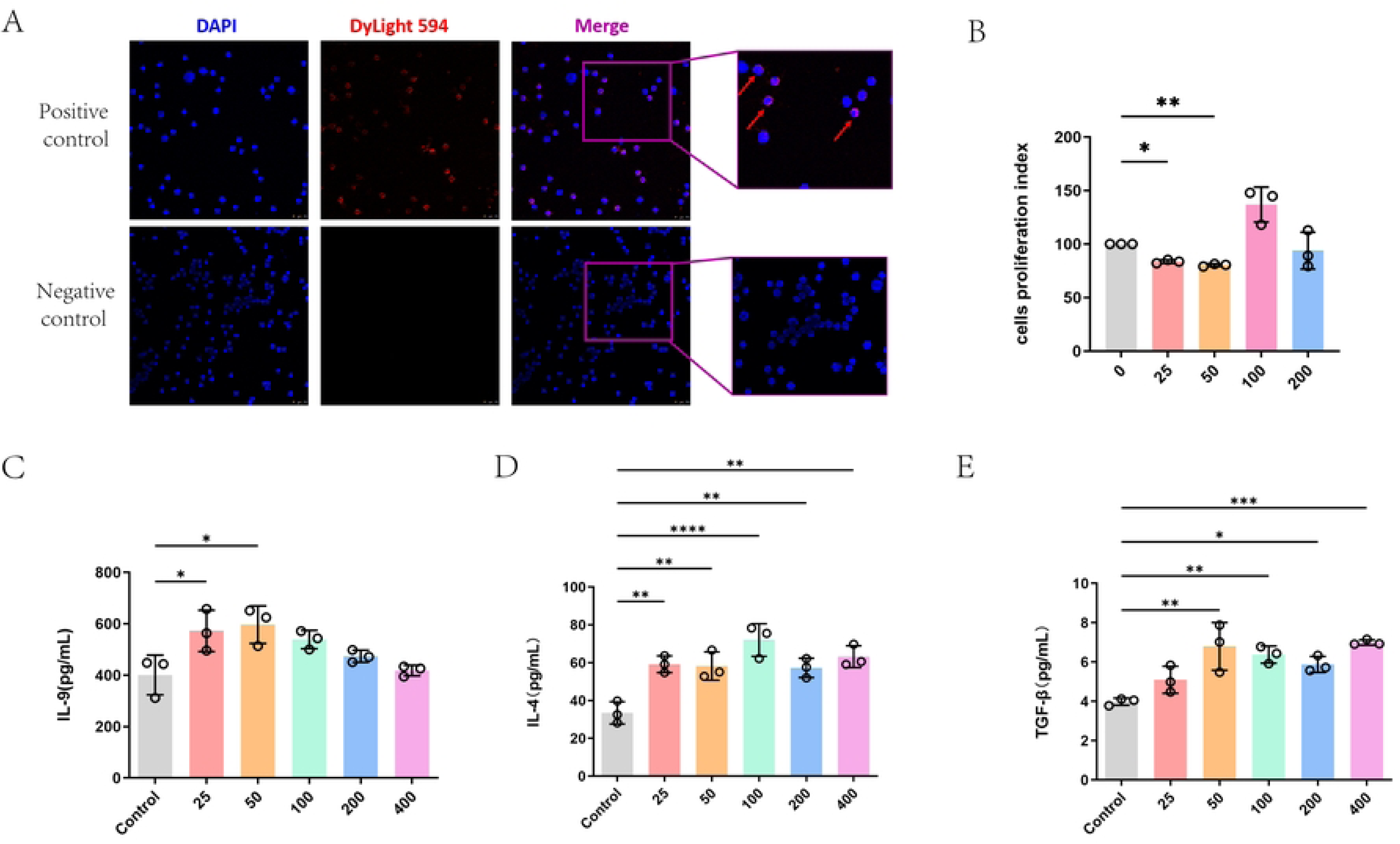
*E.granulosus* PSCs Inhibit Lymphocyte Proliferation and Modulate IL-9, IL-4, TGF-β Concentration-Dependently: Mouse splenic lymphocytes were co-incubated with PSCs for 2 h and the images of DAPI (blue) and PSCs (red) immunofluorescence co-staining were observed (A). The scale bar is 100 μ m. The effect of PSCs at concentrations of 0, 25, 50, 100 and 200 μg/mL on the viability of mouse splenic lymphocytes was detected using the CCK-8 kit (B). ELISA kits were used to detect the effects of 0, 25, 50, 100 and 200 μg/mL PSCs on IL-9 (C),IL-4(D) and TGF-β(E).

The results of cytokine detection showed that PSCs had a promoting effect on the expression of IL-9 at low concentrations, and the promoting effect was weakened at high concentrations (Fig 1C). The expression of IL-4 was significantly promoted at moderate concentrations, but was weakened but still present at high concentrations (Fig 1D). The expression of TGF-β showed a gradually enhanced promoting effect with the increase of concentration (Fig 1E).

### Identification of genes differentially expressed in the immune cells after PSCs stimulation

The DESeq software was utilized for the analysis of differential gene expression between the control group and the PSCs group. DEGs were classified as significant based on the parameters |fold change| > 2 and P < 0.05. A total of 1840 DEGs were identified. The volcano plot can visually show the overall distribution of differential genes in the two groups of samples (Fig 2A).

**Fig 2:**
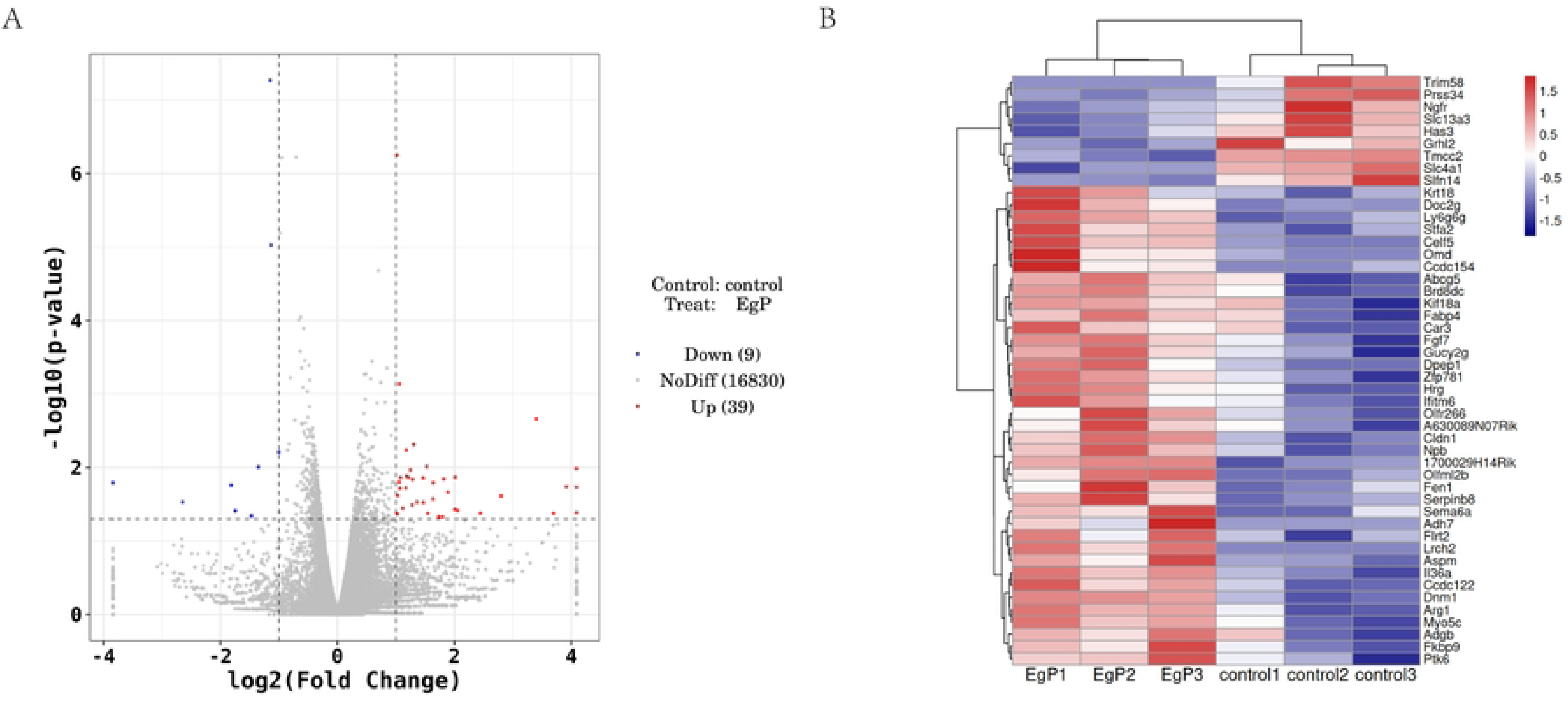
Identification of genes differentially expressed in the immune cells after PSCs stimulation: Volcano plot (A) illustrates the differential gene expression between the Control group and the PSCs-treated group. The x-axis represents the log2 Fold Change in gene expression, while the y-axis denotes the -log10 (p-value). Red dots signify up-regulated genes, blue dots represent down-regulated genes, and gray dots correspond to genes with no significant difference in expression. Heatmap (B) presents the results of hierarchical clustering analysis of gene expression across different samples. The x-axis indicates sample groups, including the PSCs-treated and Control groups, whereas the y-axis lists the names of differentially expressed genes. Color bars reflect relative levels of gene expression, with red indicating high expression and blue denoting low expression.

To evaluate the regulation of host immune cells by PSCs in vitro, we performed a heatmap with the top 20 genes most variably expressed among samples (Fig 2B). Furthermore, the mRNA expression profiles of the control group and the PSCs treatment group exhibited significant clustering within their respective groups. These clustering results suggest that PSCs has a substantial impact on the mRNA expression of various genes following immune cell stimulation, potentially influencing the regulation of host immune cell response.

### Gene Ontology and KEGG pathway analysis of differentially expressed genes after PSCs stimulation

KEGG pathway enrichment analysis of DEGs identified 20 significantly upregulated pathways (Fig 3A) and 9 downregulated pathways (Fig 3B), with comprehensive functional annotations detailed in Table 1. The up-regulated pathways were mapped to five KEGG subsystems, including environmental information processing (mmu02010), genetic information processing (mmu03450, mmu03410 and mmu03030), organismal systems (mmu04975, mmu04979, mmu04923 and mmu04961), metabolism (mmu00910, mmu00220, mmu00350, mmu00620, mmu00071, mmu00330, mmu00010, mmu00982 and mmu00980) and human diseases (mmu05218 and mmu05100). In line with these, the down-regulated pathways were aslo mapped to four KEGG subsystems, including Cellular Processes (mmu04215), organismal systems (mmu04966 and mmu04722), environmental information processing (mmu04015, mmu04014, mmu04060, mmu04010 and mmu04151) and human diseases (mmu05202). GO enrichment analysis of DEGs across biological processes (BP), cellular components (CC), and molecular functions (MF) revealed the top 30 enriched terms (Fig 3C: upregulated; Fig 3D: downregulated). Notably, DEGs were prominently linked to immunometabolic regulation and intestinal barrier integrity. These findings indicate that *E. granulosus* PSCs drive functional remodeling of host immune cells through coordinated immunometabolic reprogramming.

**Fig 3.**
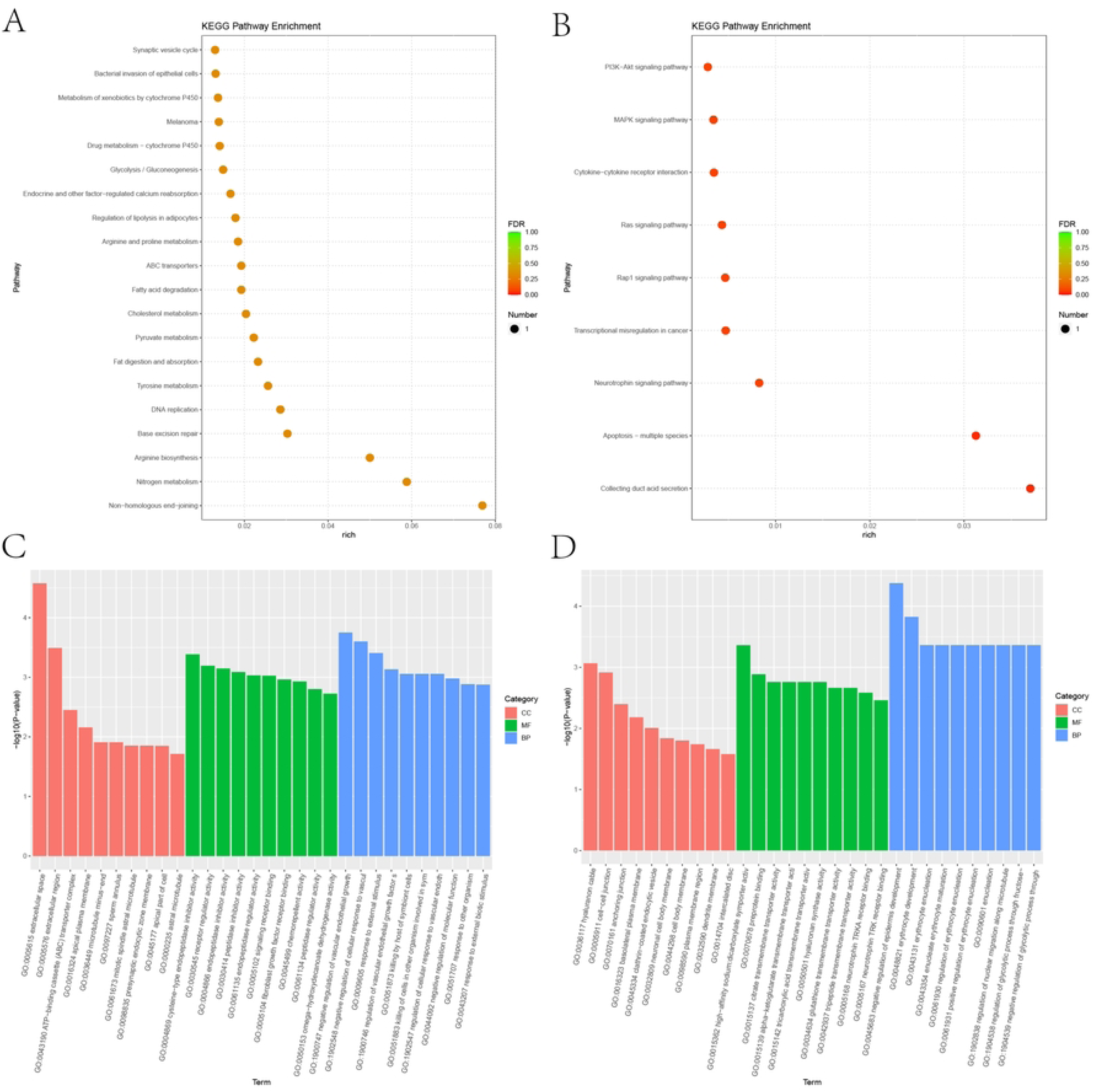
Gene Ontology and KEGG pathway analysis of differentially expressed genes after PSCs stimulation: Panel A illustrates the top 20 up-regulated pathways significantly enriched with DEGs, while Panel B depicts the top 9 down-regulated pathways associated with down-regulated DEGs, The horizontal axis denotes the enriched pathways, the vertical axis indicates the degree of enrichment, the color of the bubbles reflects the false discovery rate (FDR) value, and the size of the bubbles corresponds to the number of genes. Panels C (up-regulated) and D (down-regulated) present the GO enrichment analysis results for DEGs. In these panels, the horizontal axis represents the degree of enrichment, the vertical axis signifies the enriched functional categories, and the bars in different colors denote distinct functional categories (red for cellular component, green for molecular function, and blue for biological process).

### Screening of differential metabolites after PSCs stimulation

Integrated LC-MS/GC-MS metabolomics profiling identified 12 differentially abundant metabolites (4 depleted, 8 elevated) between controls and PSCs-treated groups, with quantitative alterations detailed in Table 2. The difference volcano plot (Fig 4A) shows the top five metabolite names with the smallest P value. The influencing factors of metabolic pathways are shown by the bubble diagram (Fig 4B). Z-score (Fig 4C) can intuitively show the overall change trend and difference degree of the quantitative value of metabolites between the experimental group and the control group in different groups and samples. The correlation heat map of differential metabolites (Fig 4D) showed that there was a synergistic or mutually exclusive relationship between different metabolites, and the metabolites had the same change trend, indicating that the relative content changes of such metabolites were positively correlated. The opposite trend is a negative correlation.

**Fig 4:**
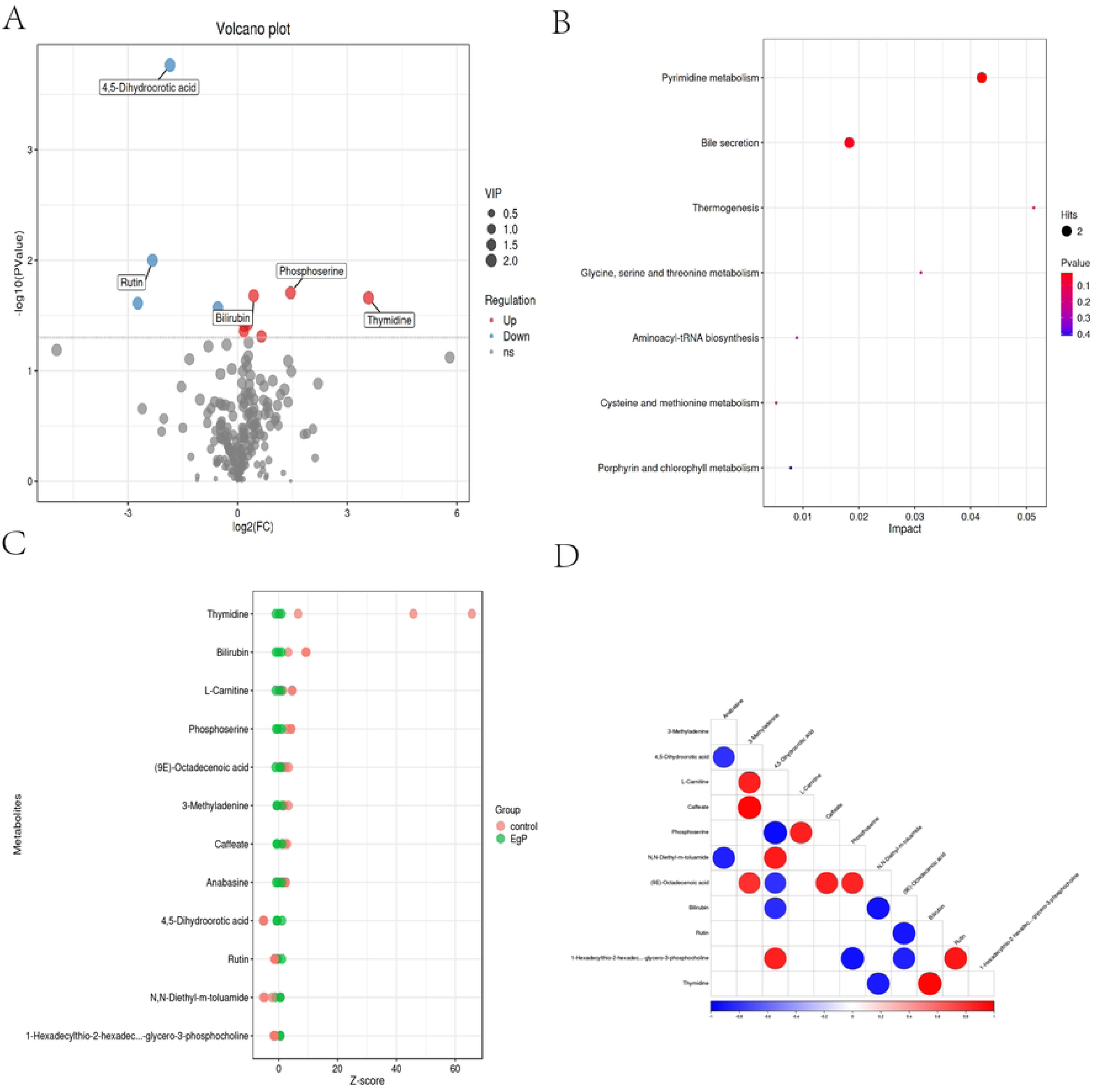
Screening of differential metabolites after PSCs stimulation: Panel A Volcano plot shows the differential expression of metabolites, the horizontal axis represents the log2 fold change of metabolites, the vertical axis represents the negative log of p value, and the color and size indicate the significance and impact value of metabolites, respectively. Panel B presents the enrichment analysis of metabolic pathways, the horizontal axis represents the influence value of the pathway, the vertical axis represents the metabolic pathway name, and the color and size represent the p-value and the number of metabolites, respectively. Panel C shows the expression of differential metabolites; the horizontal axis indicates the name of the metabolite, the vertical axis indicates the expression level of the metabolite, and the color indicates the different groups. Panel D presents the correlation network graph of metabolites, with nodes indicating metabolites and colors and connecting lines indicating correlation strength.

### Metabolomics and Transcriptomics Association Analysis

Integrated multi-omics enrichment mapping of discriminatory metabolites and mRNAs revealed both shared and unique biological pathways (Fig 5A). Bubble plot visualization quantified cross-omics concordance through pathway enrichment scores, highlighting immunometabolic axes with coordinated transcriptional-metabolomic regulation. By default, the p. Value values were ranked from small to large, and the top 20 enrichment results of differential metabolism and differential mRNA pathways were taken respectively. The nine-quadrant diagram (Fig 5B) shows the changes of chemical substances in each group in the combined analysis of transcriptomics and metabolomics. According to the difference fold threshold of mRNA and metabolites, the gray dotted line is used to divide them into 1-9 quadrants from left to right and from top to bottom, respectively. Metabolites that are consistent with or opposite to mRNA expression trends may represent genes that regulate the expression of metabolites in a positive or reverse manner. All correlation calculation results of differential mRNA and differential metabolites were selected, and the correlation clustering heatmap was drawn. The top50 mRNA and metabolism were sorted from small to large based on P.value, and the correlation heatmap was displayed (Fig 5C).

**Fig 5:**
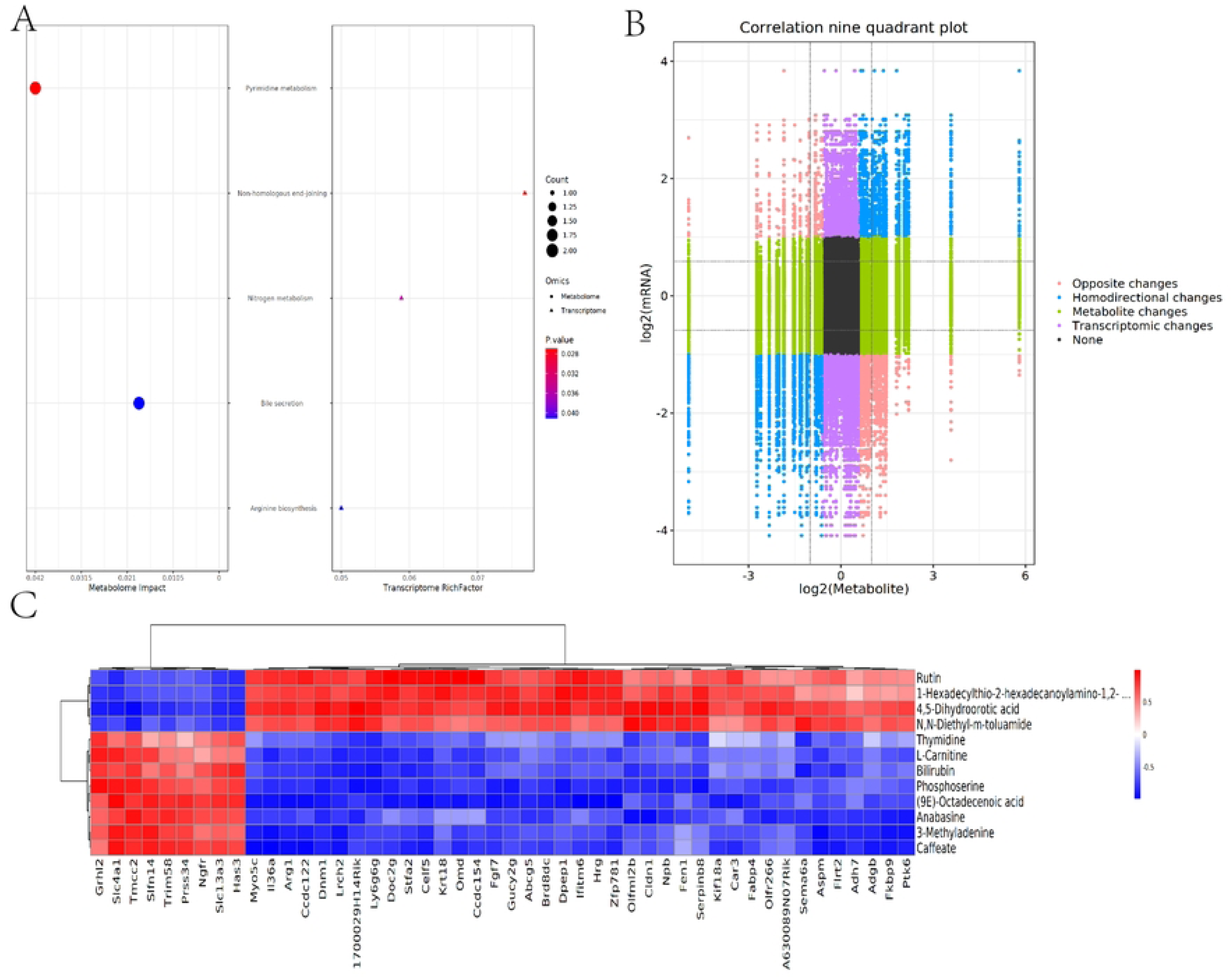
Metabolomics and Transcriptomics Association Analysis: Panel A presents the bubble plot of metabolomics impact versus transcriptome enriched factors, with the horizontal axis representing metabolomics impact values, the vertical axis representing pathway names, bubble color representing p-values, and size representing counts. Panel B is the correlation nine-quadrant plot, the horizontal axis represents the log2 fold change of metabolites, the vertical axis represents the log2 fold change of mRNA, and different colors indicate different change relationships. Panel C is a heat map showing the correlation between gene and metabolite expression, with the horizontal axis representing the gene, the vertical axis representing the metabolite, and the color representing the strength of the correlation.

## Discussion

*E. granulosus* has evolved highly sophisticated strategies to evade host immune defenses and ensure long-term survival within the host. These mechanisms can be categorized into antigenic mimicry, antigen depletion, antigenic variation, and immune modulation, all of which contribute to its persistence in the host environment [21]. *E. granulosus* secretes a diverse array of molecules that suppress host inflammatory responses. For instance, *E. granulosus* Antigen B (EgAgB) modulates the host’s inflammatory response by inhibiting the pro-inflammatory M1 macrophage phenotype and promoting the polarization of anti-inflammatory M2 macrophages [22]. This multi-omics investigation characterized the transcriptomic and metabolic responses of murine splenic lymphocytes to PSCs stimulation. RNA-Seq-derived top 20 DEGs were prioritized for functional validation to delineate parasite-driven immunomodulatory mechanisms. The majority of these DEGs were associated with enzyme activity (hydrolysis, transfer, isomerism), receptor signaling (growth factors, G-protein-coupled receptors), and gene expression regulation. Additionally, chemical types such as substance transport (ions, metabolites), cell structure/movement (keratin, myosin), transcriptional regulation (transcription factors, RNA-binding proteins), and immune modulation indicated that DEGs play critical roles in cellular metabolism, development, and immune responses. These findings corroborate earlier biochemical studies on *E. granulosus*, which demonstrated that cyst fluid contains heat shock proteins, cytoskeletal proteins, and other molecules that potentially mediate biological processes and host-parasite interactions [23, 24].

In this study, following the stimulation of mouse immune cells with protoscolex crude antigen, the up-regulated pathways were identified within five subsystems of the KEGG database: Upregulated pathways were primarily associated with five KEGG functional categories: environmental information processing, genetic information processing, metabolism, organismal systems, and human diseases. Conversely, downregulated pathways clustered within four subsystems: cellular processes combined with three shared categories (environmental information processing, organismal systems, and human diseases). The upregulated “Synaptic vesicle cycle” pathway may be associated with neural signaling and synaptic function [25], indicating that PSCs could potentially modulate the immune response by influencing neurotransmitter release. The down-regulated “PI3K-Akt signaling pathway” may impact cell proliferation, apoptosis, and metabolism [26], aligning with the regulatory effects of PSCs on host cell functionality. The upregulated entries for “plasma membrane” and “cell adhesion molecule binding” suggest that PSCs might regulate intercellular interactions and signaling by altering the composition and function of the cell membrane. Conversely, the down-regulated entries for “cell surface” and “signal transducer activity” imply that PSCs may reduce the capacity of cells to respond to external stimuli, which could be associated with the immunosuppressive effects induced by PSCs.

Immunometabolism constitutes a frontier research domain that elucidates the complex interplay between metabolic pathways and host immunity, as well as its implications for overall health. Previous studies have shown that echinococcus infection modulates the expression of genes associated with various metabolic pathways, including cancer-related pathways, fat digestion and absorption, and retinol metabolism[27]. Consistent with these findings, this study identified the downregulation of Rhein and Rutin, alongside the upregulation of Bilirubin, Phosphoserine, and Thymidine in the experimental group. Moreover, significantly enriched pathways such as “Pyrimidine metabolism,” “Bile secretion,” and “Thermogenesis” collectively suggest that the host reprograms its metabolic network to coordinate immune defense and energy supply. The accumulation of Thymidine is closely linked to the activation of pyrimidine metabolism, which may promote immune cell proliferation or tissue repair [28]. Meanwhile, Phosphoserine contributes to nucleotide synthesis through serine metabolism [29]. The upregulation of Bilirubin exhibits dual functions: antioxidation by scavenging free radicals (compensating for the reduced levels of Rutin) and immunosuppression by inhibiting T cell activation [29], potentially fostering an immune-tolerant microenvironment conducive to parasite survival. Given the limited capacity of lipid synthesis in *E. granulosus*, essential lipids must be acquired from the host [30]. Elevated levels of L-Carnitine and the activation of the “Thermogenesis” pathway indicate enhanced fatty acid oxidation to meet increased energy demands [31, 32]. Hosts deploy multidimensional defenses via mechanisms involving pyrimidine metabolism, bile regulation, and thermogenic pathways, while parasites may exhibit adaptive metabolic strategies, such as Bilirubin-mediated immunomodulation.

In this study, we demonstrated that the protein of *E. granulosus* PSCs regulates the function of mouse splenic lymphocytes through perinuclear localization. The concentration-dependent inhibition of proliferation indicates that low concentrations of PSCs may induce cytotoxicity, while high concentrations attenuate inhibition or correlate with adaptive cell survival mechanisms. The transient activation of IL-9 by PSCs and the sustained upregulation of IL-4/TGF-β suggest that PSCs may promote immune shifts through differential regulation of Th9-type cytokines. Notably, the dose-dependent increase in TGF-β levels highlights the central role of PSCs in mediating immunosuppression. By integrating the results of KEGG and GO enrichment analyses, we speculate that PSCs induces functional reprogramming of host immune cells by modulating specific pathways and gene functions. This reprogramming likely involves multiple biological processes, including metabolic adjustments, signaling alterations, and immune response regulation. For instance, metabolic-related pathways such as “Glycolysis and Gluconeogenesis” may regulate energy metabolism, thereby influencing immune cell activity. Additionally, modulation of signaling pathways like the “PI3K-Akt signaling pathway” by PSCs may alter cellular responses to stimuli, affecting the intensity and duration of immune responses. Furthermore, PSCs may regulate the interaction between intestinal microbiota and the host by influencing genes and pathways associated with intestinal barrier function, which is crucial for maintaining intestinal homeostasis and preventing inflammatory diseases. Metabolomic profiling delineated distinct metabolite signatures, and integrated multi-omics data elucidated parasitic protein-driven immunomodulatory mechanisms—particularly splenic lymphocyte polarization dynamics and evolutionarily conserved host defense strategies against helminthic invasion.

## Acknowledgments

We thank the State Key Laboratory of Pathogenesis, Prevention and Treatment of High Incidence Diseases in Central Asia for providing experimental platforms. We would also like to thank all those who contributed to this work.

## Data Availability

All relevant data are within the paper and its Supporting information files.

## Table

Table1: The differential up-regulated and down-regulated pathways.

Table2: Extreme significant difference metabolites.

